# Revealing nanoscale structure and interfaces of protein and polymer condensates via cryo-electron microscopy

**DOI:** 10.1101/2024.02.15.577400

**Authors:** Aoon Rizvi, Bruna Favetta, Nora Jaber, Yun-Kyung Lee, Jennifer Jiang, Nehal S. Idris, Benjamin S. Schuster, Wei Dai, Joseph P. Patterson

## Abstract

Liquid-liquid phase separation (LLPS) is a ubiquitous demixing phenomenon observed in various molecular solutions, including in polymer and protein solutions. Demixing of solutions results in condensed, phase separated droplets which exhibit a range of liquid-like properties driven by transient intermolecular interactions. Understanding the organization within these condensates is crucial for deciphering their material properties and functions. This study explores the distinct nanoscale networks and interfaces in the condensate samples using a modified cryo-electron microscopy (cryo-EM) method. The method involves initiating condensate formation on electron microscopy grids to control droplet size and stage in the phase separation process. The versatility of this method is demonstrated by imaging three different classes of condensates. We further investigate the condensate structures using cryo-electron tomography which provides 3D reconstructions, uncovering porous internal structures, unique core-shell morphologies, and inhomogeneities within the nanoscale organization of protein condensates. Comparison with dry-state transmission electron microscopy emphasizes the importance of preserving the hydrated structure of condensates for accurate structural analysis. We correlate the internal structure of protein condensates with their amino acid sequences and material properties by performing viscosity measurements that support that more viscous condensates exhibit denser internal assemblies. Our findings contribute to a comprehensive understanding of nanoscale condensate structure and its material properties. Our approach here provides a versatile tool for exploring various phase-separated systems and their nanoscale structures for future studies.

## INTRODUCTION

Liquid-liquid phase separation (LLPS) is a process in which a homogenous solution separates into two or more liquid phases.^1^ This phenomenon is observed in various molecular solutions including small molecules,^2^ polymers,^3^ peptides,^4^ and proteins.^5^ LLPS of proteins is commonly observed in living systems and plays an important role in transcription, DNA repair, chromatin organization, and disease formation.^6–8^ LLPS of synthetic polymers plays an important role in chemical synthesis, development of biomimetic systems, and also the food industry.^9^ Consistent across these different cases is the generation of liquid-like droplets composed of high concentration of material immersed within a dilute solution.

Condensed material that results from phase separation is often characterized as dynamic, and droplets can undergo fusion and exchange material with their surroundings.^10^ This liquid-like nature of droplets arises from the transient nature of intermolecular interactions which hold the condensates together, including hydrogen bonds, hydrophobic interactions, and electrostatic interactions.^6^ Intermolecular interactions are known to drive the self-assembly of synthetic and biological macromolecules into materials with distinct nanostructures.^11^ For example, synthetic polymers can assemble into particles with lamellar and bicontinuous microstructures.^12^ In biological systems, lipids organize into bilayers to form cell membranes or lipid droplets.^13,14^ However, the structures that emerge within phase separated systems are largely unexplored due to the difficulty in characterizing condensate material on the nanoscale. There is some evidence that protein-based and polymer based condensates can form nanoscale structures, such as *in silico* work showing the emergence of structured liquids with some combinations of charge patterned disordered proteins and polymers.^15^ Additionally, recent cryo-electron tomography (cryo-ET) based studies of phase-separating folded proteins support that liquid-like condensates can exhibit a range of nanoscale structures ranging from amorphous to ordered morphologies.^16^

It is important to explore the existence of an internal nanostructure of condensates because these may be connected to their material properties and function. The same intermolecular interactions that drive the formation of condensed material may contribute to its material properties. The function of condensates depends in part on their material properties,^17^ because these influence internal molecular dynamics, the partitioning of non-phase separating macromolecules, and interactions with other assemblies or structures.^18^ Therefore, the connection between condensate nanostructure, material properties, and function should be explored.^19^ Recent work has made progress towards quantitatively describing the material properties of condensates.^20^ Interestingly, some of these findings suggest that material properties are not homogenous within condensates, suggesting a heterogeneous organization of the condensed material on the nanoscale.^21,22^ In the case of naturally occurring protein-based condensates, a close connection between the viscosity of a condensate and its function has been identified.^23,24^ Condensates are often found to be highly dynamic when operating as a reaction center, while rigidity can be important for functions such as storage, protection, or organization of components.^25^ Better understanding the nanoscale organization within condensates may help us understand their function and how they function.

Several techniques have been applied to elucidate the nanoscale structure of condensates, including super-resolution microscopy,^26^ small-angle X-ray scattering (SAXS),^27^ and cryo-electron microscopy methods (cryo-EM).^28^ Cryo-EM is emerging as a powerful method for elucidating nanoscale condensate morphology because of its nanometer resolution and retention of the native-like structure of a specimen. Recently, Zhang et. al. studied tetra-nucleosome condensate forming mechanism using cryo-electron tomography (cryo-ET) and specialized electron microscopy grids with a streptavidin layer for high specimen loading. Their data revealed a new mechanism of nucleation and growth of condensates.^29^ Additionally, Mahamid and co-workers recently developed a method for cryo-EM of reconstituted condensates.^28^ In their protocol, condensates are allowed to form on grids for several minutes prior to vitrification of the specimen. Furthermore, they utilize correlative light microscopy as a first step to explore the preformed condensate specimens prior to electron microscopy imaging. Their electron microscopy observations revealed network-like structures and clusters of biomacromolecules within the formed condensates. While these methods are impactful by enabling nanoscale elucidation of condensate structures, the requirement of special grids and additional imaging methods limits the broad application of these approaches.

As the field of polymer and biopolymer phase separation continues to grow, methodology that is easily adaptable by non-experts is required to expand the exploration of condensates at the nanoscale. Here we present a workflow for preparing cryo-EM samples of three different classes of condensates by modifying routine cryo-EM sample preparation. We focus on protein condensates to explore different nanoscale features using our approach. We observe internal networks within condensates and a distinct interface which resembles a membrane visually. We also reconstructed the 3D volume of protein condensates using cryo-ET and relate the 3D condensate structure to their material properties. The 3D analysis also revealed inhomogeneities, such as a distinct core and shell within the nanostructure of condensates.

## Results & Discussion

### Method Development

The most common method for preparation of cryoEM samples is the blotting plunge freezing method developed by Dubochet and coworkers.^30,31^ In this method, a small volume of sample (1 – 4 µL) is deposited on a grid surface, followed by blotting by a filter paper that removes a majority of the deposited solution, leaving a thin film of sample that is then frozen. The challenge of applying this method to condensates is that condensates often quickly grow into droplets > 1 micron in diameter, which are too thick to resolve using high-resolution electron microscopy. To address this issue, we devised a method to initiate condensate formation on the grid which enables droplets to be vitrified at smaller sizes. We take advantage of factors that determine protein phase separation such as concentration, temperature, and salt concentration to induce condensation on the grid. This procedure is shown in Figure 1 for protein condensates that formed upon reduction of salt concentration via mixing of a low salt buffer but can be adapted for any type of condensation initiation (see Supporting Information). First, a known volume of buffer is applied to the grid. After the buffer is applied, protein solution is added to the grid and the solution becomes turbid, indicating the formation of condensate droplets. We found that addition of the diluent buffer prior to the protein sample minimized protein sticking to the cryo-EM grids. The excess solution is blotted away for 3 seconds, and the sample is plunged. By including waiting period between the steps described above, condensate growth can be controlled for the desired droplet size and stages in the phase separation process. As shown in Figure 1, we tested waiting periods between steps 2 and 3 or steps 3 and 4, such that condensates are allowed to grow either before or after the blotting process. We found that if the imaging goal is to capture early-stage formation, then it is best to allow condensate growth following step 2. This is because any evaporation effects will be minimal at this stage, and the formation of droplets can be observed as the liquid on the grid becomes turbid. Sometimes, macromolecules like proteins and polymers can undergo shear induced morphological transitions upon blotting during the sample preparation for cryo-EM (Supplementary Information).^32,33^ If the protein or polymer condensates are susceptible to shear induced morphological transitions, it is best to let the condensates grow following the blotting step. This will allow the condensates to reach equilibrium again.

**Figure 1:**
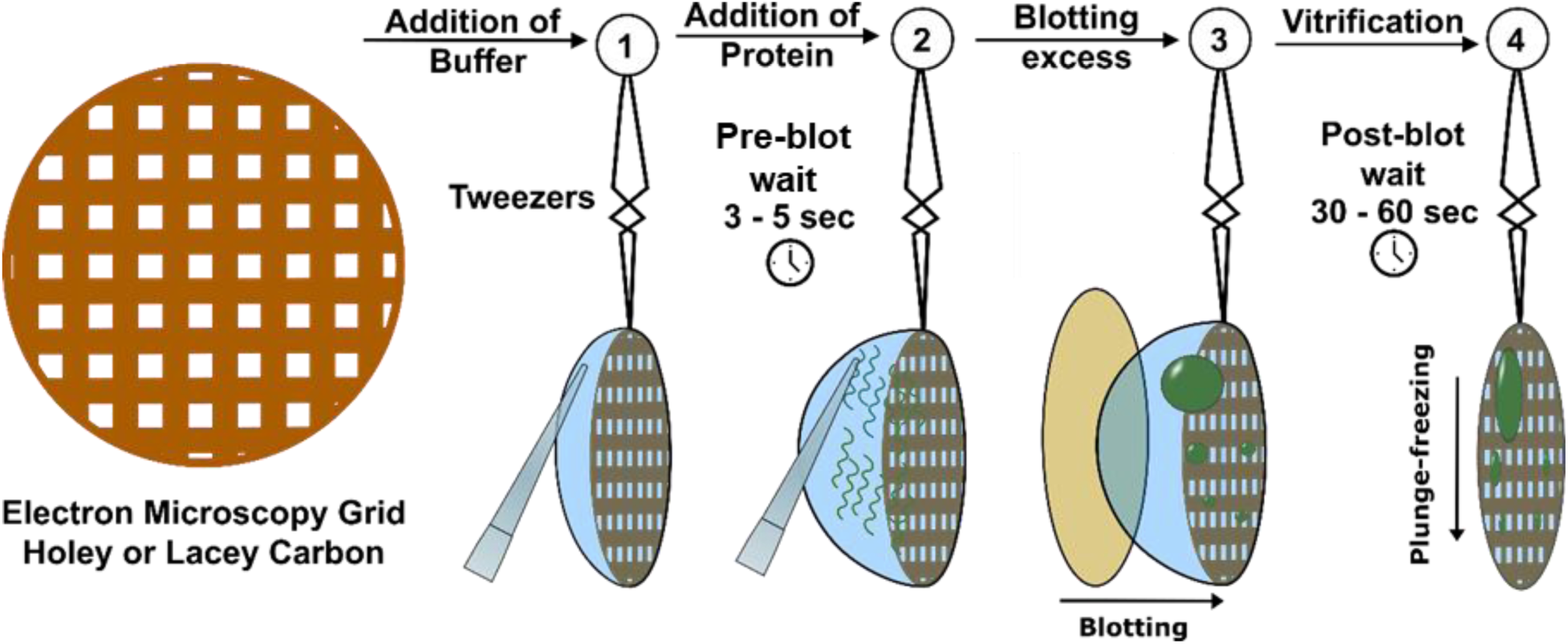
Overview of on-grid sample preparation for cryo-EM samples of RGG condensates (See SI for different condensate preparations). In this procedure, phase separation was induced by reducing the salt concentration of the protein solution. In Step 1, we add a known amount of diluent to the grid. In Step 2, the protein solution is added, and droplet formation is initiated. Up to 5 seconds of wait time was used prior to moving on to the next step. This ensures the two solutions mix well and condensates are formed. In Step 3, excess solvent is blotted away for 3 seconds, resulting in a thin layer of the specimen. Up to 60 seconds of wait time was allowed prior to moving on to the next step to minimize any shear-induced artefacts from blotting. In Step 4, the grid is plunged in a cryogen bath to vitrify the thin layer.

To demonstrate the versatility of this technique, we applied it to three classes of condensates, including complex coacervates,^34^ protein condensates,^35^ and non-ionic block copolymer coacervates.^36^ For complex coacervates, we used the combination of single stranded DNA(66 nt) and poly-Lysine (PLK100). Similar complex coacervates of DNA and PLK have been used as model systems to understand membranelles organelles^37,38^ and to test coacervates for nucleotide delivery. For the protein samples we focused on the RGG domain from LAF-1, a prototypical arginine/glycine-rich intrinsically disordered protein (IDP) involved in P granule assembly in *C. elegans* (see Supplementary Figure 1). LAF-1 RGG contains amino acids that can promote several modes of interactions, including electrostatic, π–π, and cation-π interactions.^39,40^ The patterning of amino acids within IDPs has a direct relationship to the phase behavior and material properties of condensates.^41^ We explored two versions of this RGG domain, the wild type RGG domain (WT) and a variant in which the amino acids were shuffled (SH) to create blocks of oppositely charged residues, to visualize how sequence patterning may affect the nanoscale organization of material within the condensates. SH RGG was first designed by Schuster, Dignon, and coworkers (labeled “RGGshuf-press”) when studying the effects of sequence perturbations on protein phase behavior.^41^ The SH variant has an abundance of anionic residues in the first half of the sequence, and an abundance of cationic residues in the second half, in contrast with the WT sequence, which has a relatively even distribution of cationic and anionic residues throughout. Lastly, for non-ionic block copolymer coacervates we used polyethylene oxide-*block-*poly methyl methacrylate in mixtures of dioxane and water.^36^

Figure 2 shows representative images from the three classes condensates that were imaged (see Supplementary Figures 2-5 for more examples). We found that results were consistent across multiple preparations of the samples. Typical cryo-EM specimens are vitrified onto grids with a perforated carbon layer. The preferred location for imaging is over the holes to minimize interference of the carbon layer; each image shown in Figure 2 was taken over a hole in the grid. In our study, condensates were present both inside of the hole and on the carbon layers. The condensates are easily identified at nominally lower magnifications as they display good contrast against the ice layer. This allows easy screening of the specimen prior to higher magnification imaging. Each class of condensates displays nanoscale internal features suggesting that this is a common feature of macromolecular liquid droplets. The protein and complex coacervates display liquid-like smooth curved structures and have a distinct interface with the dilute solution, whereas the block copolymer coacervates appear to have rougher interfaces and larger internal features. This is likely due to the microphase separation of the two incompatible blocks, which gives it a porous structure.^15^

**Figure 2:**
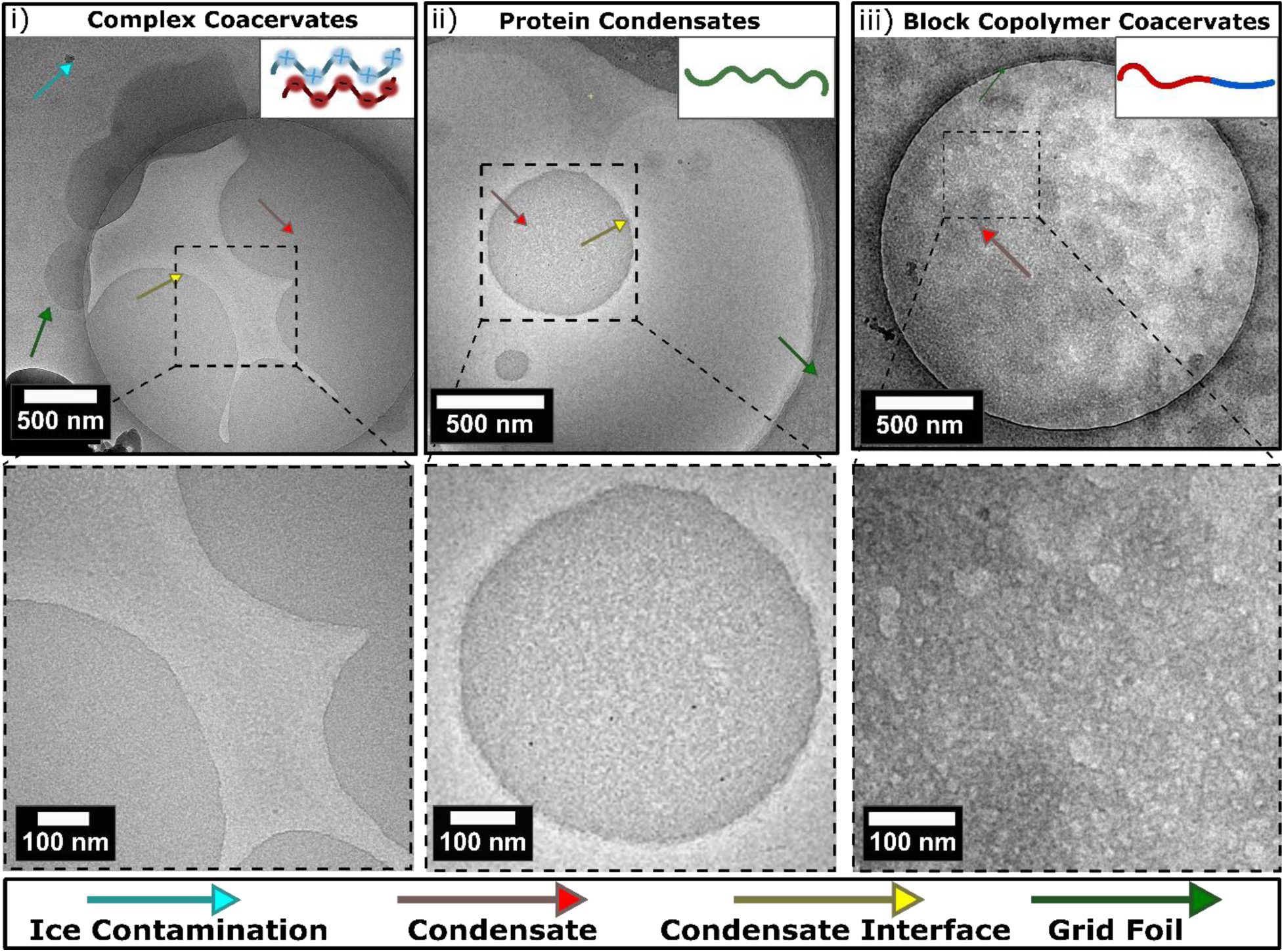
Cryo-EM images of a variety of condensates/coacervates. i) Complex coacervates made of 5 mg mL^-1^ poly-Lysine (cationic) and 5 mg mL^-1^ single-stranded DNA (anionic) (Representative image from n = 50 images taken). The curved interface of a large condensate can be seen here in the center of the grid hole, grid holes are large circles as seen in (i,ii&iii) without any substrate surface. These areas are ideal for imaging to minimize noise from substrate surfaces (pointed out by green arrows). The bottom image shows a close-up of the boxed area. ii) Protein condensate of 100 µM WT RGG in 150 mM NaCl, 20mM Tris-HCl buffer (Representative image from n > 100 images taken). Two spherical condensates can be seen within the center of the grid hole. The bottom image shows a close-up of larger condensate. iii) Block copolymer coacervates of 10 mg mL^-1^ polyethylene glycol-*block*-polymethyl methacrylate polymer (non-ionic) in 1:4 dioxane:water mixture (Representative image from n > 100 images taken). The bottom image shows a close-up of porous structured condensate material. The bright regions represent the hydrated hydrophilic block, and the darker regions represent the collapsed hydrophobic block; the two regions appear to be in a continuous morphology.

### Comparison with Dry State TEM

While cryo-EM is an ideal electron microscopy method for imaging solvated structures, conventional dry-state TEM is still commonly used.^42^ It is well known that the drying of macromolecular assemblies changes their organization as they are held together by weak noncovalent interactions and the solvent is an inherent part of their interactions and structure.^42,43^ Figure 3 compares the structures of the block copolymer and protein condensates using both cryoEM and dry-state TEM. In Figure 3i, the dried coacervates resemble solid particles with no preservation of the porous condensate structure, as seen in Figure 3ii. Similarly, in Figure 3iii dried condensates composed of WT RGG can be seen with no structure or shape as compared to the spherical droplet with internal structure shown in Figure 3iv. The same trend can be seen for SH RGG condensate in Supplementary Figure 6. While dry-state TEM may be a more accessible method, these data stress the importance of preserving the hydrated structure of coacervates for understanding their nanoscale structure.

**Figure 3:**
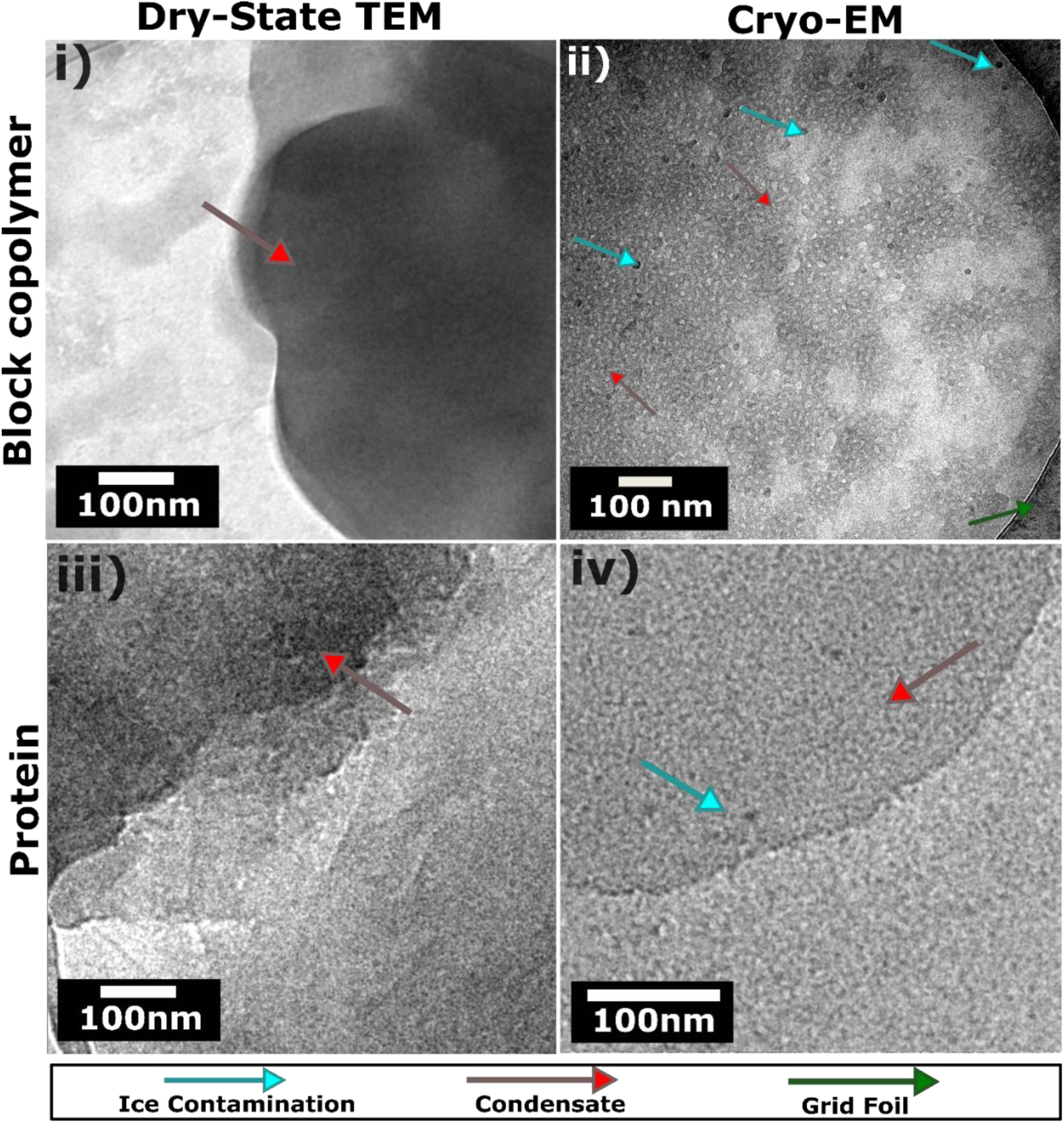
Comparison of Cryo-EM and dry-state TEM of condensates. i) Dry-state TEM of block copolymer coacervates. (Representative image from n = 20 images taken) Dried condensate material can be seen with no preserved structure. ii) Cryo-EM image of block copolymer coacervates with the preserved internal organization. (Representative image from n = 50 images taken) iii) Dry-state TEM of WT RGG condensate. Dried condensate material can be seen with no preserved structure. (Representative image from n = 20 images taken) iv) Cryo-EM image of WT RGG condensate showing internal network structure with a distinct interface. (Representative image from n > 100 images taken) Dry-state specimens are prepared on lacey carbon grids, which appear as a web-like structure in the background of i & iii.

### Condensate Interface Nanostructure

Biological condensates are commonly referred to as membraneless organelles (MLOs), as these droplets are found in cells with no lipid membranes.^44^ While membrane-bound organelles utilize the lipids at their interface to control the transport of proteins and other biomolecules, MLOs have to achieve a similar control through other mechanisms.^44^ Recent studies have highlighted the importance of condensate interfaces as sites for enhanced reaction efficiencies and partitioning of biomolecules.^45–47^ However, little is known about the structure and morphology of biomolecular condensate interfaces.

In our data, we observe a distinct interface between the dilute and protein-dense phase, which qualitatively resembles a membrane (Figure 4i-v). Inspecting 100 individual condensates for both RGG variants, WT and SH, we observed that over 95% of the condensates showed a distinct interface (Figure 4 vi). As seen in Figure 4, the interface boundary can be seen as a darker band which wraps the spherical droplet across the whole interface (Figure 4i). This darker region correlates to an electron dense area as compared to its surrounding, suggesting a higher concentration of the protein at the interface and/or accumulation of salts at the interface. Recent computational studies aimed at understanding protein conformations at the condensate interface suggest several ways in which proteins might organize themselves at this boundary.^48,49^ Measuring the interface of the protein-based condensates revealed the thickness of the surface layer to be ∼6 nm (Figure 4ii). Interestingly, this surface layer appears to be longer than the expected RGG domain length of ∼3.5 nm inside of condensates based on previous simulation work.^50^ One possible explanation of our observation is that the proteins organize at the interface and stretch out along their long axis. This is supportive of the work by Farag et al. in which Monte Carlo based simulations suggested that proteins at the interface of condensates organize perpendicular to the interface of the droplet as shown in Figure 4iii (top), resulting in a highly stretched confirmation which is distinct from the conformation of proteins in the core of a droplet.^48^ In contrast, it is possible that the protein chains are collapsed and tightly packed together and that multiple layers form the distinct interface of 6nm thickness. This is supportive of the work by Wang et al. in which they also used Monte Carlo based simulations to study protein conformations at condensate interfaces.^49^ They found that proteins at the interface are mostly collapsed except the tail ends that stick out of the condensate to minimize entropy loss as shown in Figure 4iii (bottom). In both cases, we would expect the condensate interface to appear as we observe in our data. The compacted interface also aligns with the recent findings of Zhang et al. which showed tetranucleosomes are more compacted at condensate interface via cryo-ET.^29^ Yet another factor that may be contributing to the unique interface we observe is the accumulation of salt ions (NaCl) at the interface, neutralizing local protein charge.^46^ Future studies using correlative cryo-EM and FRET or other methods will be required to further understand the condensate interface.^51^

**Figure 4:**
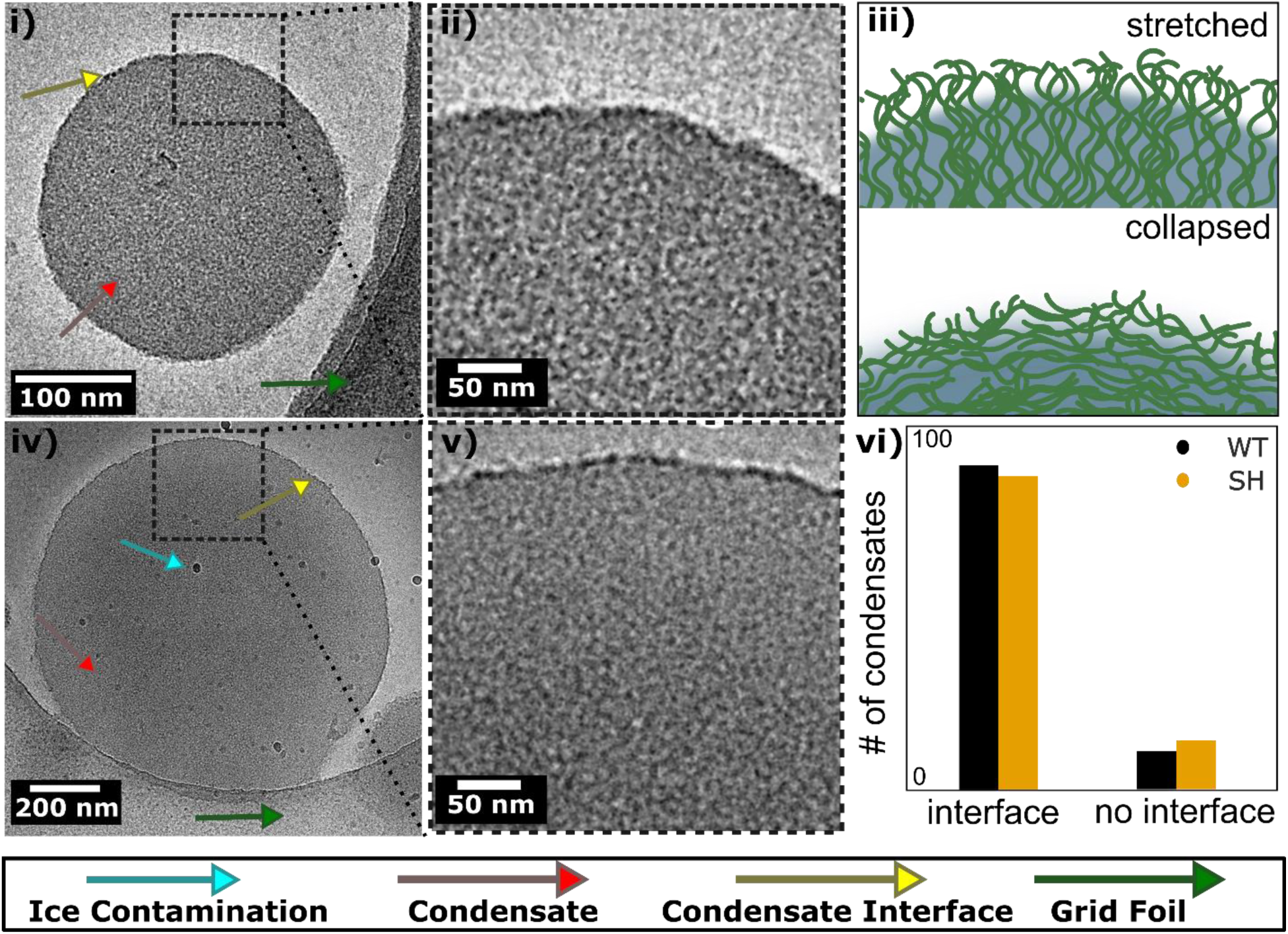
Interfaces of the of WT and SH RGG condensates. i) WT RGG condensate. (Representative image from n > 100 images taken) ii) Crop of WT RGG condensate in which the distinct interface can be seen between the condensed and dilute phases. iii) Schematics of possible protein organization at condensate interface, (top) perpendicular and stretched organization of the protein, (bottom) collapsed organization of the protein with only the chain ends being perpendicular to the interface. iv) SH RGG condensate. (Representative image from n > 100 images taken) v) Crop of SH RGG condensate in which the distinct interface can be seen between the condensed and dilute phases. vi) Number of condensates with clear interface structure for both RGG variants, WT and SH.

In less than 5% of the condensate inspected (n > 100), we observe inhomogeneities at the condensate interfaces. As seen in Supplementary Figure 7i, the interface is broken at the location pointed with a white arrow. In Supplementary Figure 7ii the spherical condensate displays a patchy interface, and Supplementary Figure 7iii shows an example with no distinct interface but a gradient edge. These anomalies could be a result of the blotting process cryo-EM sample preparation. Lastly, we also found several examples of nanoscale fiber formation adjacent to the interfaces of RGG WT condensates, specifically at necks that suggest locations of condensate breakage, as shown in Supplementary Figure 8. Although RGG-based condensates are not prone to fibrillation, the shear forces that exist during sample preparation may also be contributing to their formation (Supplementary Figure 9).^52,53^ Although an investigation of the specific structures we observed at the interface is beyond the scope of this work, our images provide additional data to the current body of work investigating protein organization at the condensate interface.^48,49^

### Internal Condensate Nanostructure

Next, we investigated what could be learned from Cryo-EM about the internal structure of condensates. Recent work has pointed to condensates as being porous and inhomogeneous materials,^54,55^ however, direct evidence of these structural properties is limited. To investigate the internal nanostructure of condensates, we turned to electron tomography. In this method, we collect a series of EM images of a single droplet at multiple angles (-60° to + 60°). The series of 2D projection images can then be reconstructed to recover the 3D volume.^56,57^ For tomographic analysis the sample preparation is similar to the protocol described above, but with the addition of exogenous fiducial markers, typically gold nanoparticles, that help with downstream data analysis and 3D reconstruction. Since mechanical stage drift and beam induced sample motion during experimental data acquisition occur,^58,59^ fiducial markers are used to facilitate alignment of the images acquired at different tilt angles.

Figure 5 depicts the 3D condensate structure of WT RGG and the SH RGG variant. Generally, condensates appear as a porous material composed of a network of protein (electron dense material) and tunnels, likely filled with buffer (brighter, less electron dense regions). For example, the average of the central slices of WT and SH RGG condensates shown in Figure 5i and 5iv clearly shows network formation within the condensate. Taking an intensity profile of a selected region shows the varying electron density within the condensates, with peaks representing protein densities and troughs representing pores or voids (Figure 5v). The presence of such networks with voids has been predicted recently using Monte Carlo simulations of IDPs.^48^ It was suggested that these voids emerge due to the regions of high vs. low crosslinking densities for a given protein.

**Figure 5:**
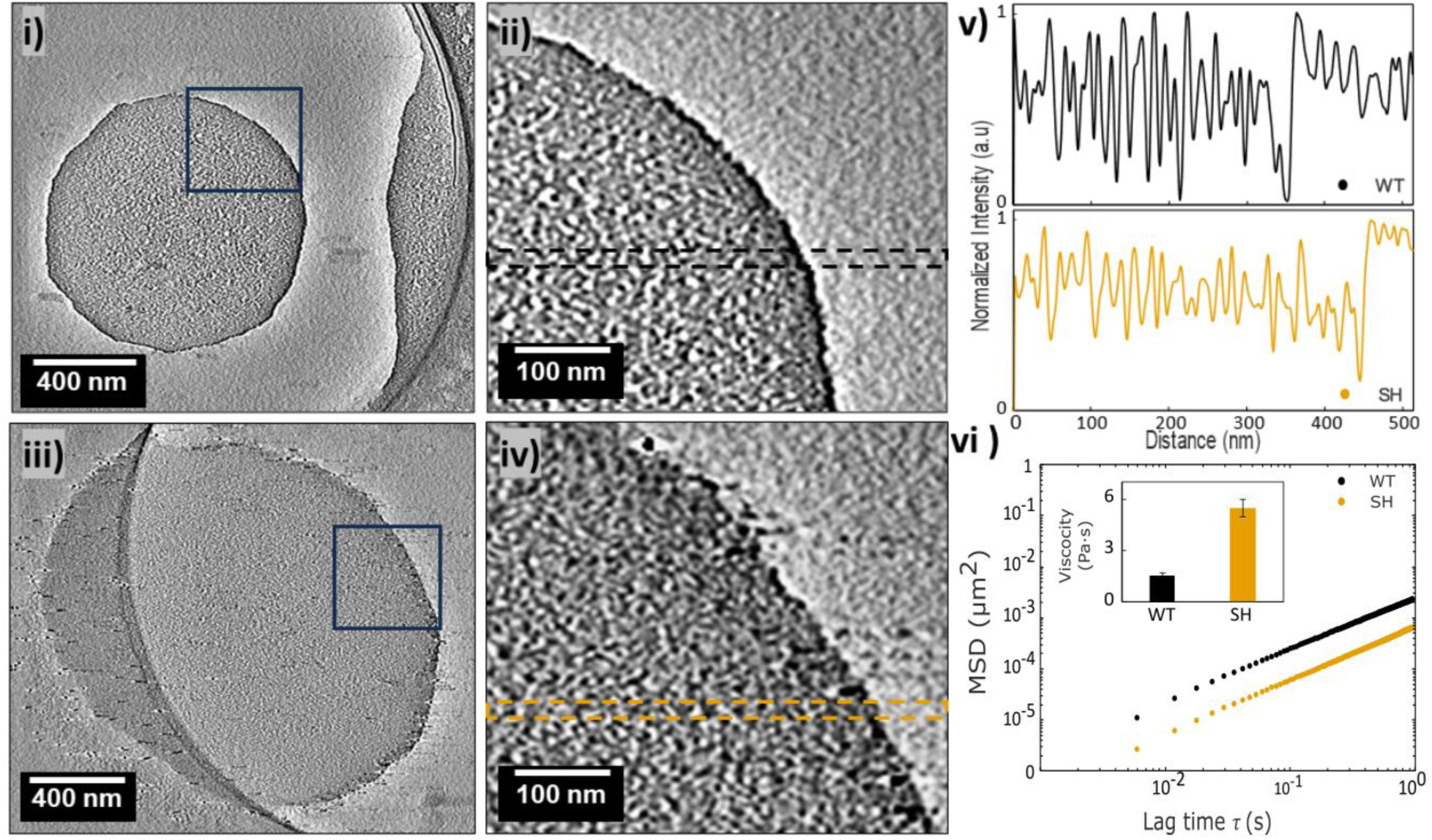
Comparison between the internal condensate structure and material properties of WT RGG and SH RGG condensates. i) Tomogram of WT RGG condensate. ii) Zoomed in section of tomogram (i) showcasing porous internal morphology and a distinct interface of WT condensate. iii) Tomogram of SH RGG condensate. iv) Zoomed in section of tomogram (iii) showcasing dense internal morphology and a distinct interface of SH condensate. v) Intensity profiles from WT and SH condensates of the boxed areas as shown in (ii) and (iv) respectively. vi) Ensemble mean-squared displacement versus lag time for WT and SH RGG condensate, inset shows the viscosity calculated from the particle tracking results.

Using tomographic analysis, we can explore the effects of charge patterning on condensate nanostructure. The charge patterning of an IDP has been identified as a key determinant of its phase separation behavior,^41,61^ since it contributes to the strength of intermolecular interactions that can occur. However, little is known about how charge patterning affects the nanostructure and function of condensates. For example, recently, it was discovered that charge patterning of IDPs can control partitioning of transcriptional machinery within condensates of the Mediator complex IDP, thus affecting its function.^62^ Still, we lack a structural understanding of how this mechanism works. We suspect that one contributing factor which determines partitioning behavior inside a condensate may be a change in condensate structure that happens because of IDP charge patterning.

In Figure 5i and 5iii, a difference in structure within the condensates can be seen between the RGG WT and SH condensates. WT condensates appear to have larger voids and pores, while SH condensates appear to have a compact network structure. These organizational differences on the scale of a few nanometers may arise due to the difference in charge patterning between SH and WT condensates, and this structural difference may dictate partitioning behaviors as seen in recent literature.^62^ Further studies combining labeled particles and cryo-EM may elucidate how condensate nanoscale structure may dictate partitioning within biomolecular condensates.

Based on the difference in nanoscale structure between the WT and SH RGG condensates, we hypothesized that the two condensates may also have different material properties. To test this, we measured the viscosity of the two condensates using video particle-tracking micro-rheology (VPT). VPT allows us to measure condensate viscosity by tracking the displacement of beads inside of condensates. We embed 500 nm fluorescent particles into the condensates and track their movement over time. Although the 500 nm beads used in VPT are much larger than the nanostructures we are observing, their Brownian motion is sensitive to the bulk rheology that arises from the nanostructures. As seen in Figure 5vi, the slope of the mean square displacement (MSD) indicates that both condensates behave as viscous fluids.^63^ However, the SH condensates are around 3 times more viscous than their WT counterparts (Figure 5vi). This agrees with our nanoscale observations in which SH condensates appear to be denser and with less pores compared to the WT. The two condensates in our study show that differences in the nanoscale structure of a condensate can have small effects on its material properties like, measured viscosity (Figure 5vi). Interestingly, the viscosities of condensates can vary multiple orders of magnitude, which is explained by varying strengths of molecular interactions.^64^ However, the relationship between the possible viscosities that emerge from the nanoscale organization is still widely unexplored. While research is being conducted on the sequence determinants of material properties of condensates,^65,66^ our data and methodology adds to the understanding of the structural determinants that influence condensate material properties and provides a way to explore the varying nanoscale structure of condensates.

Lastly, our tomographic analysis reveals a unique core-shell morphology of RGG WT condensates. Here, we define core-shell morphology as a condensate containing lower density of material at its core compared to at the surface layer. As shown in Figure 6i-ii, voids seen as the bright spots within the condensate are both larger and more numerous towards the center of condensates. Closer to the interface, the condensate material is distributed more evenly and densely without the presence of many voids. Through the 3D isosurface view shown in Figure 6ii-iii, we can verify that the shell appears to have a higher density of protein compared to the core (the shell region is highlighted in Figure 6ii by the yellow arcs). These results suggest that RGG condensates have a porous, void filled core and a denser layer of condensate material surrounding the core. Core-shell condensates have been observed previously, typically made up of two or more macromolecular species and fall into the larger category of “multiphase condensates”.^47,67–69^ Our observation is different as only one protein type creates the core-shell structure at the nano-scale. This difference in morphology of the condensate material in the core and surface should manifest into varying material properties within the same condensate. In fact, such differences have been encountered before in condensates: using single particle tracking, the viscosity of TFEB condensates was found to be higher towards the surface than the interior of droplets.^70^ Importantly, the method we utilized to measure viscosity of condensed material would not be sensitive to these differences in material properties, given our probes are 500 nm in size, similar to the diameter of the entire condensates observed using cryo-EM. How this core shell morphology of condensates scales with condensate size should be examined using other techniques. Overall, our results add to the growing evidence that condensates can be heterogenous materials and that caution is necessary when measuring their material properties. The existence of this type of architecture in other protein-based condensates should be further explored to advance our understanding of structure-property relationships in biomolecular condensates.

**Figure 6:**
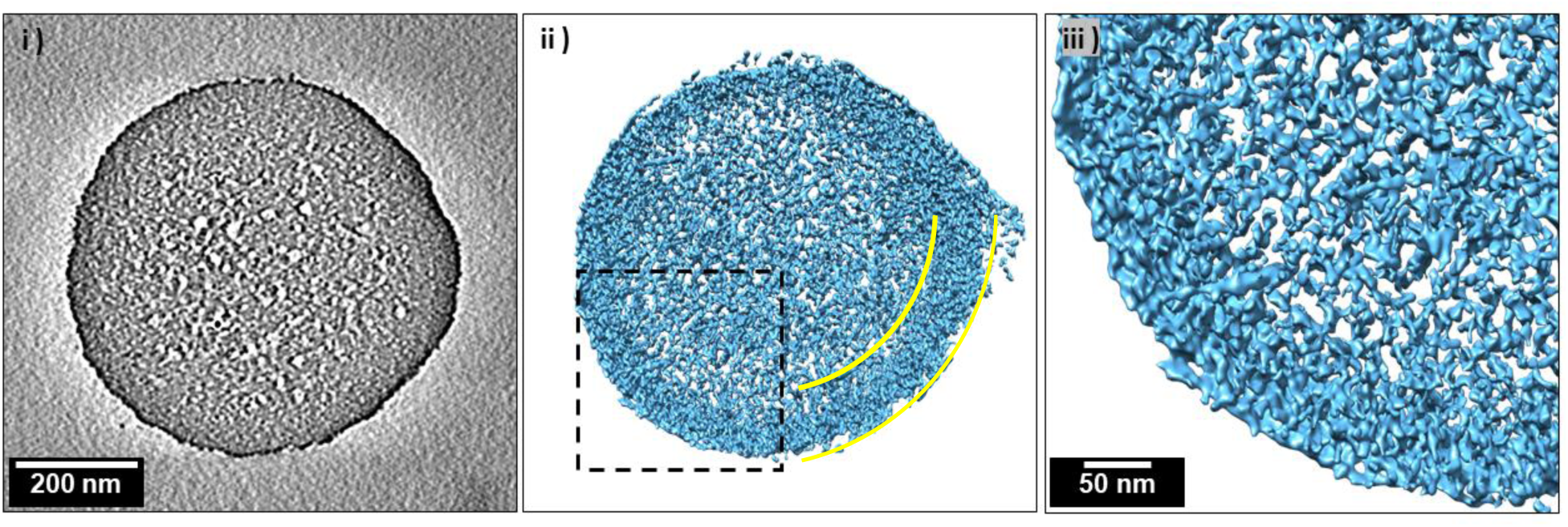
Visualization of core-shell morphology and voids within WT RGG condensate observed using cryo-ET. i) Tomogram of RGG WT showcasing a dense shell of condensate material around a porous core. ii) Isosurface view of the tomogram (i) shows the pores visible only within the core of the condensate. Yellow arcs are used to highlight the shell of dense condensate material. iii) Zoomed-in square area from (ii) helps to see the dense shell layer of condensate material towards the edge of the condensate.

## Conclusion

In conclusion, this study revealed how proteins are organized within biomolecular condensates and at their interface using a modified cryo-EM approach. Our approach, applied across three classes of condensates, demonstrates its versatility and potential to significantly advance our understanding of phase-separated systems. While we mostly focused on protein condensates in this report, we envision this methodology to be adopted widely to study nanoscale structure of several kinds of phase separated systems, like block copolymer coacervates, complex coacervates, and others. The insights gained from these experiments can help lead to a more complete understanding of condensate structures and their functions. We discovered unique condensate internal and interface structures, which align with *in silico* and in vitro work of others. We also show how the internal structure of the condensates is related to their viscous properties. Furthermore, we observe core-shell morphology of protein condensates using cryo-ET, which could explain the observations of heterogenous properties of condensates. With the ability to image condensates at the nanoscale in a simple manner, future studies can be carried out to explore the large set of parameters that affect condensate formation, their properties, and how these relate to biological function.

## AUTHOR INFORMATION

**Notes:** The authors declare no competing financial interest.

## ACKNOWLEDGMENTS

The authors would like to acknowledge Prof. Alex Marraas for providing complex coacervates used in this study. The authors acknowledge Prof. Jeetain Mittal for helpful discussions regarding condensate interfaces. A.R. was supported by the Rowland Fellowship at UC Irvine Chemistry. The authors acknowledge the use of facilities and instrumentation at the UC Irvine Materials Research Institute (IMRI), which is supported in part by the National Science Foundation through the UC Irvine Materials Research Science and Engineering Center (DMR-2011967). B.F. and B.S.S. acknowledge funding from the National Institutes of Health (R35GM142903). N. J. was supported by the NSF award MCB-2046180. The authors acknowledge the use of the Rutgers CryoEM & Nanoimaging Facility for cryo-ET data collection and would like to thank Jason Kaelber and Emre Firlar for their support.

## Data availability statement

Tomograms of WT and SH RGG tomograms have been deposited in the EMDataBank under accession codes EMD-43500, EMD-43502, EMD-43503, and EMD-43504.

## ASSOCIATED CONTENT

## Supplementary Information

Additional information is supplied as Supporting Information. Correspondence and requests for materials should be addressed to J.P.P.

