## Supplemental Information for "Revealing nanoscale structure and interfaces of protein and polymer condensates via cryo-electron microscopy"

### Materials and Instruments:

### Proteins:

Throughout this paper, WT RGG denotes the RGG domain from LAF-1 (residues 1-168 of LAF-1) and SH RGG denotes the charged shuffled version.[1,2] All proteins included a hexahistidine tag at the C-terminus for immobilized metal affinity chromatography.

**RGG WT**
MESNQSNNGGSGNAALNRGGRYVPPHLRGGDGGAAAAASAGGDDRRGGAGGGGY
RRGGGNSGGGGGGGYDRGYNDNRDDRDNRGGSGGYGRDRNYEDRGYNGGGGG
GGNRGYNNNRGGGGGGYNRQDRGDGGSSNFSRGGYNNRDEGSDNRGSGRSYNN
DRRDNGGDGLEHHHHHH

**RGG SH**
MGGYGYGSSGDGGGDDYGDARYVPPHLRGYGDGAGDDGGDNNDDSDDADRDYN
GGLSGGAGGNSGGDGENGGDGNGRNNARSGNNRGGNGNYRYFGANYGAGEGRG
RNGQGGEGSGNNRGGGGRYGRRRRQGSRGGRGSGGNYGGNSNRSGRAGGRDN
NARNRRRNGSLEHHHHHH

**DNA** (Provided by Prof. Alex Marras, UT-Austin)**:**

**DNA(66)**
TCAACATCAGTCTGATAAGCTATGGATACTCGTCTGGACTACTTACTCACTCATTCA
TCACTATCT

### Block Copolymers:

Polyethylene oxide-*block*-poly methyl methacrylate ($PEO_{45}$-*b*-$PMMA_{300}$) was synthesized as previously described.[3] The solid polymer was dissolved in dioxane shortly prior to preparing cryo-EM samples to minimize concentration variation.

### Protein expression and purification

For bacterial expression, plasmids were transformed into BL21(DE3) competent *E. coli* (New England BioLabs). Colonies picked from fresh plates were grown for 8 h at 37 °C in 1 mL LB + 1% glucose while shaking at 250 rpm. This starter culture (0.5 mL) was then used to inoculate 0.5 L cultures. For RGG WT and SH proteins, cultures were grown in 2 L baffled flasks in autoinduction medium (Formedium) supplemented with 4 g/L glycerol at 37 °C overnight while shaking at 250 rpm. The pET vectors used contained a kanamycin resistance gene; kanamycin was used at concentrations of 50 μg/mL in cultures [4]. After overnight expression at 37 °C, bacterial cells were pelleted by centrifugation at 4100 x g at 18 ˚C. Pellets were resuspended in lysis buffer (1 M NaCl, 20 mM Tris, 20 mM imidazole, Roche EDTA-free protease inhibitor, pH 7.5) and lysed by sonication. Lysate was clarified by centrifugation at 25000 x g for 30 minutes at 25 ˚C. The clarified lysate was then filtered with a 0.22 μm filter. Lysis was conducted on ice, but other steps were conducted at room temperature to prevent phase separation.

Proteins were purified using an AKTA Pure FPLC with 1 mL nickel-charged HisTrap columns (Cytiva) for affinity chromatography of the His-tagged proteins. After injecting proteins onto the column, the column was washed with 1 M NaCl, 20 mM Tris, 20 mM imidazole, pH 7.5. Proteins were eluted with a linear gradient up to 1 M NaCl, 20 mM Tris, 500 mM imidazole, pH 7.5. Proteins were dialyzed overnight at 42°C using 7 kDa MWCO membranes (Slide-A-Lyzer G2, Thermo Fisher) into physiological buffer (150 mM NaCl, 20 mM Tris, pH 7.5) or high salt buffer (300 mM NaCl, 20 mM Tris, pH 7.5). Proteins were snap frozen in liquid $N_2$ in single-use aliquots and stored at -80 °C.

For microscopy experiments, protein samples were prepared as follows: RGG protein aliquots were thawed above the phase transition temperature and diluted into the desired concentration with 20 mM Tris, 150 mM NaCl, pH 7.5 buffer, or diluted with 20 mM Tris, 0 mM NaCl, pH 7.5 buffer if lower salt was desired. Protein concentrations were measured based on their absorbance at 280 nm using a Nanodrop spectrophotometer (ThermoFisher); RGG WT and SH proteins were mixed in a 1:1 ratio with 8 M urea to prevent phase separation during concentration measurements.

**SDS-PAGE**

For chromatographically purified proteins, SDS-PAGE was run using NuPAGE 4-12% Bis-Tris gels (Invitrogen) and stained using a Coomassie stain (SimplyBlue SafeStain; Invitrogen).

**Cryo-EM and Cryo-ET**

All Cryo-EM experiments were performed on a JEOL 2100 equipped with a 200 keV field emission gun and a OneView camera at the Irvine Materials Research Institute, University of California, Irvine. All Cryo-ET experiments were performed on Talos Artica cryo-electron microscope (Thermo Fisher Scientific) equipped with a post-column BioQuantum energy filter and K2 direct electron detector (Gatan, Inc) at Rutgers University, Institute for Quantitative Biomedicine.

**Sample Preparation for Cryo-EM and Cryo-ET**

For all samples 200/400 mesh copper grids with 1-2 um holes were used. Each grid was glow discharged prior to sample preparation. For Cryo-ET samples, 6 nm gold nanoparticles (Thermo Fisher) were added onto each grid as fiducial markers.

**Protein Condensates:**

Protein condensates of RGG WT and its charge variant RGG SH were prepared by changing the salt concentration below which each protein undergoes phase separation. To capture the condensates onto the cryo-EM grids, phase separation was carried out on each sample grid inside the humidity chamber. First, a known volume of buffer is applied to the grid. Note: It is better to add a buffer which will dilute the overall sample first, in order to minimize proteins sticking to the cryo-EM grids. After the buffer is applied, protein solution is added to the grid and the solution becomes turbid, indicating the formation of condensate droplets. After waiting between 0-5 seconds of pre-blot time, the excess solution is blotted away for 3 s and the sample is plunged immediately. For cryoET samples of RGG WT and SH the samples were blotted after 30s to 1 min of pre-blot wait time. This wait time did not show a significant difference in the samples when imaged.

**Block Copolymer Coacervates:**

Block copolymer coacervates form in our system by addition of water into a dissolved solution of the polymers.[3] $PEO_{45}$-*b*-$PMMA_{300}$ 10 mg $mL^{-1}$ in dioxane undergoes LLPS at 25% $H_2O$. To capture the coacervate droplets onto the cryo-EM grid, we prepare our solution of polymers in dioxane and apply 3 µL onto the grid while the grid is in the humidity chamber (99% RH). The sample is the blotted immediately for 3 s and plunged with no post blot time. Since dioxane is hydroscopic, the solution absorbs water and increases water content while in the humidity chamber. This way the droplets are formed and trapped onto the TEM grid. Preforming coacervates results in large coacervates due to their fast kinetics, this creates very thick ice layers inhibiting cryo-EM imaging.

**Complex Coacervates:**

Complex coacervates were composed of 66 nt ssDNA (negatively charged) and poly-L-Lysine ($PLK_{100}$) (positively charged) prepared as described previously.[5] To capture coacervate droplets onto the cryo-EM grids, we induced phase separation onto the grid inside the humidity chamber. Phase separation was induced by mixing the two solutions of each macromolecule; the concentrations were adjusted to balance out the positive and negative charges to maximize droplet formation.[6] To a glow discharged grid we added DNA solution (1.5 μL) followed by $PLK_{100}$ solution (1.5 μL). The solution turned turbid indicating the progress of the phase separation process. The sample was then blotted for 3 s and plunged immediately with no post-blot time.

*Grid Preparation for Cryo-ET:*

The LAF-1 RGG WT and SH condensates were initially kept at 44 °C to prevent phase separation. They were individually mixed with 6 nm gold nanoparticles as fiducial markers to facilitate tilt series alignment during image processing. An aliquot of 3.5 μL of the condensate sample was applied to glow discharged Quantifoil holey grids (R2.0/2.0, Cu, 200 mesh; Quantifoil). For vitrification, we used a Leica EM GP plunger (Leica Microsystems, Buffalo Grove, IL, USA) with a humidity (95%) and temperature (20°C) controlled chamber. Prior to vitrification, the condensate samples were allowed to phase separate on the grid in the plunger chamber for either 30 or 60 seconds before blotting for a total of 4 seconds. Plunge-frozen grids were stored in a liquid nitrogen dewar until imaging.

*Cryo-ET data collection and data processing*

Grids were loaded into a Talos Artica cryo-electron microscope (Thermo Fisher Scientific) equipped with a post-column BioQuantum energy filter and K2 direct electron detector (Gatan, Inc). Screening was done at 3,000x magnification. 2D projection images and title series of target areas were collected at 31,000x magnification with a pixel size of 4.30 Å/pixel. Image settings used were spot size 9, 70 µm objective aperture, and nominal defocus of -5 µm. The tilt series ranged from -60° to 60° at 3° step increments. The total dose was about 120e/Å2 per tilt series.

Alignment of tilt series and reconstruction of tomograms were done using the EMAN2 tomography workflow (Chen) and IMOD (Mastronarde). 3D map visualization and analysis were done using Chimera (University of California, San Francisco) (Petterson).

**Video Particle-Tracking Microrheology (VPT)**

500 nm diameter yellow-green carboxylate-modified polystyrene beads (FluoSpheres, Invitrogen) were used for VPT microrheology measurements. Each protein sample was prepared as previously described. Microrheology experiments were prepared by mixing the 200 µL protein sample with the fluorescent tracer beads before initiating droplet assembly in a 96-well plate (#1.5 high- performance cover glass, Cellvis). The samples were incubated at room temperature for 45 min and then were observed under the microscope to verify that the tracer beads were embedded in the condensates. Next, the samples in the well plate were centrifuged at 300xg for 30 seconds to form a

condensate layer or larger-size droplets (>30 µm in diameter); the purpose of this step was to avoid boundary effects and prevent flow of the condensates. Epifluorescence video imaging was initiated at the 1 hr timepoint with fluorescence excitation using a 475 nm LED (Colibri 7; Zeiss). Videos of the tracer beads diffusing within the condensate were collected at 200 frames per second for 2000 frames. Imaging was conducted at room temperature (18 – 20 °C). For each protein, two independent samples were made on different days, and 10-12 videos were collected from each sample, with each video containing ~10-50 tracer beads. The TrackPy particle tracking code was used to analyze the collected videos, starting with extracting particle trajectories. The mean squared displacement (MSD) was calculated from the trajectories of individual beads, followed by calculating the ensemble-average MSD. In general, the ensemble-average MSD often scales as a power law with lag time $\tau$, as given by the following equation:

$$MSD(\tau) = 2dD\tau^{\alpha}$$

where d is the number of dimensions (here d=2, since data collection and analysis were conducted in the x-y plane), D is the diffusion coefficient, and α is the diffusivity exponent. For a purely viscous fluid, the diffusivity exponent α is close to unity. α values for all the condensates tested were in the range of 0.9-1.05. Assuming a purely viscous fluid, with the system at equilibrium, the condensate viscosity η is then calculated using the Stokes-Einstein equation:

$$D = \frac{k_B T}{6\pi\eta R}$$

where $k_B$ is the Boltzmann constant, T is the temperature (Kelvin), and R is the tracer bead radius.

**Supplementary Figures:**

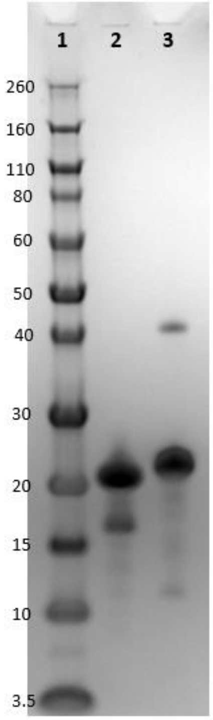


**Supplementary Figure 1:** SDS-PAGE of purified proteins expressed for this study. 1: Ladder (kDa), 2: RGG WT, 3: RGG SH

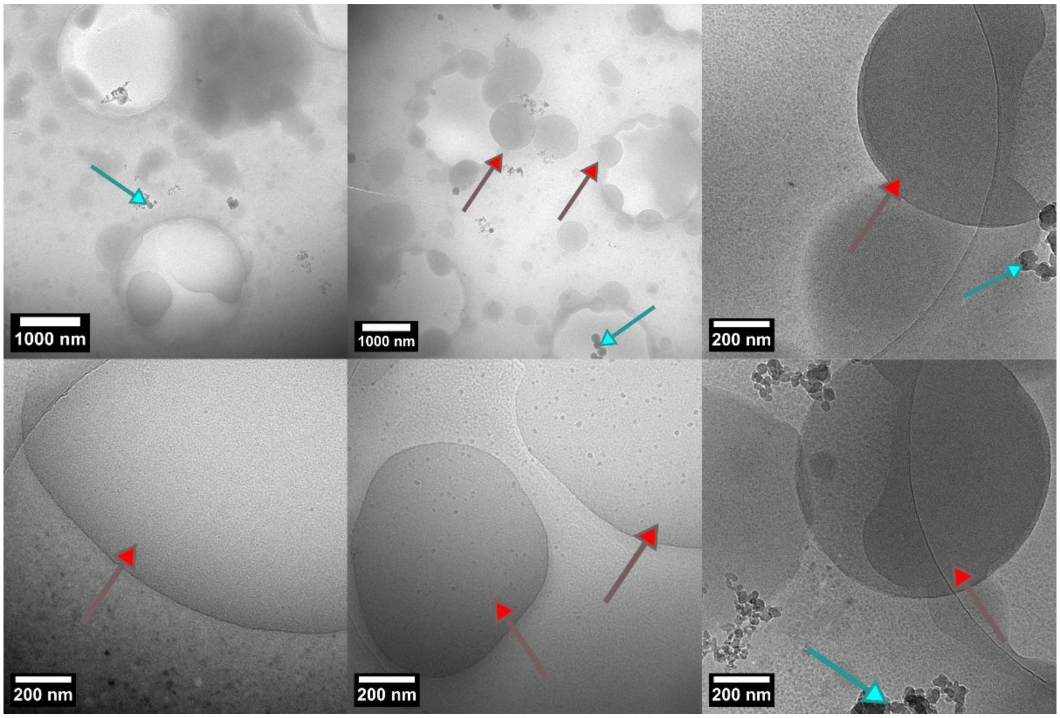


**Supplementary Figure 2**: Additional cryoEM images of complex coacervates. The varying mags showcase the size dispersity and the flattening effect. The images are taken from different samples. Blue arrows indicate ice contamination, red arrows help identify the condensates.

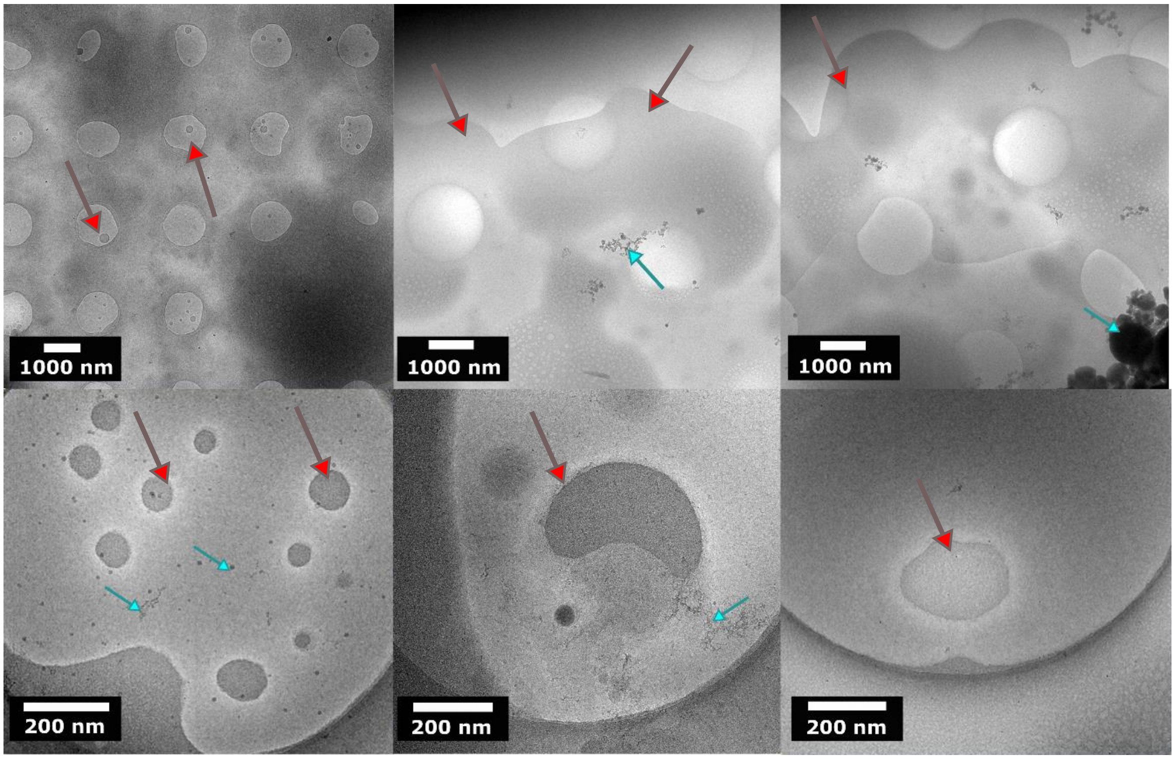


**Supplementary Figure** 3: Additional cryoEM images of WT RGG condensates. The varying mags showcase the size dispersity and the flattening effect. The images are taken from different samples. Blue arrows indicate ice contamination, red arrows help identify the condensates.

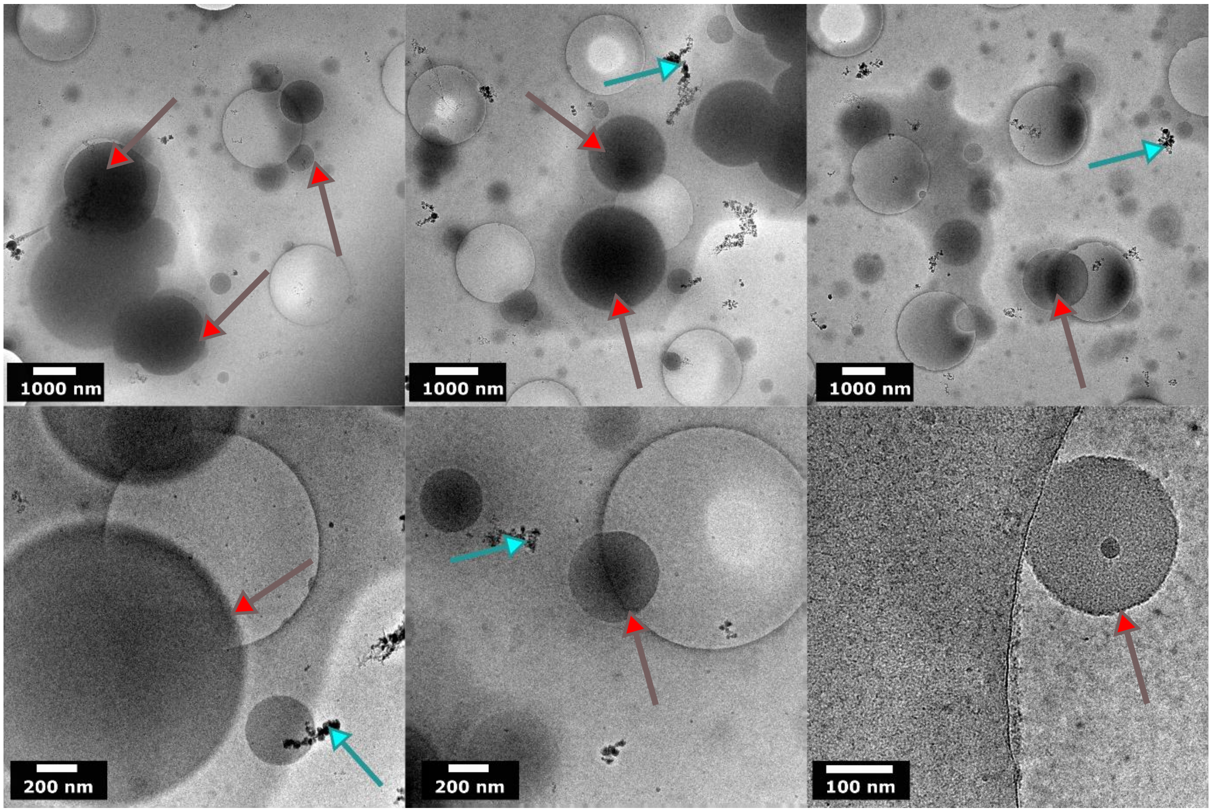

**Supplementary Figure 4:** Additional cryoEM images of SH RGG condensates. The varying mags showcase the size dispersity of the condensates. The images are taken from different samples. Blue arrows indicate ice contamination, red arrows help identify the condensates.

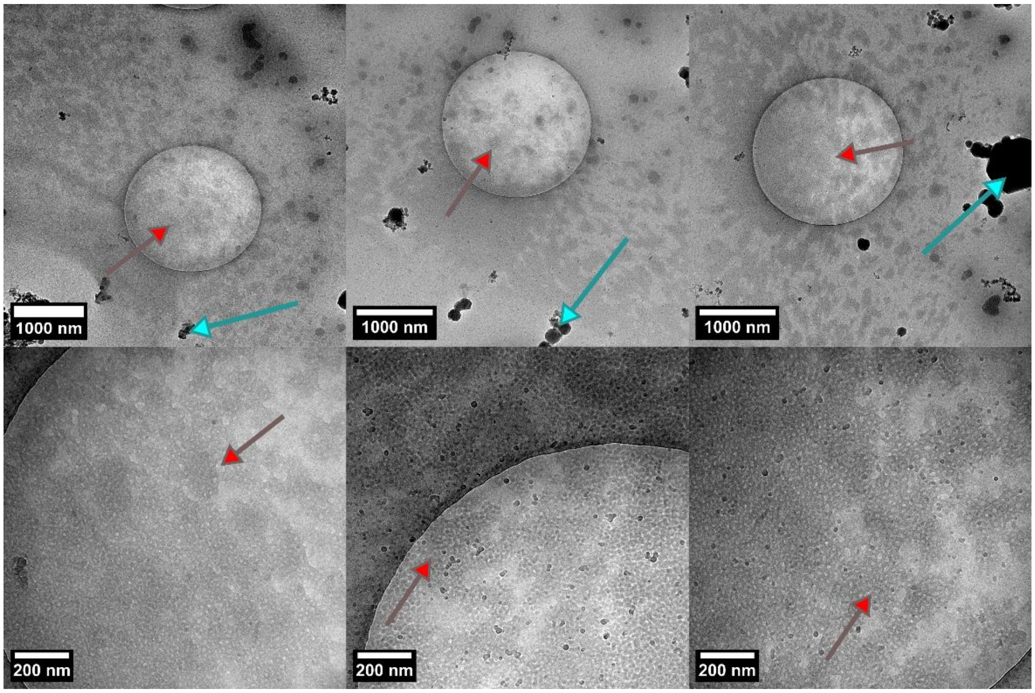


**Supplementary Figure 5**: Additional cryoEM images of block copolymer coacervates. The varying mags showcase the size dispersity and the flattening effect. The images are taken from different samples. Blue arrows indicate ice contamination, red arrows help identify the condensates.

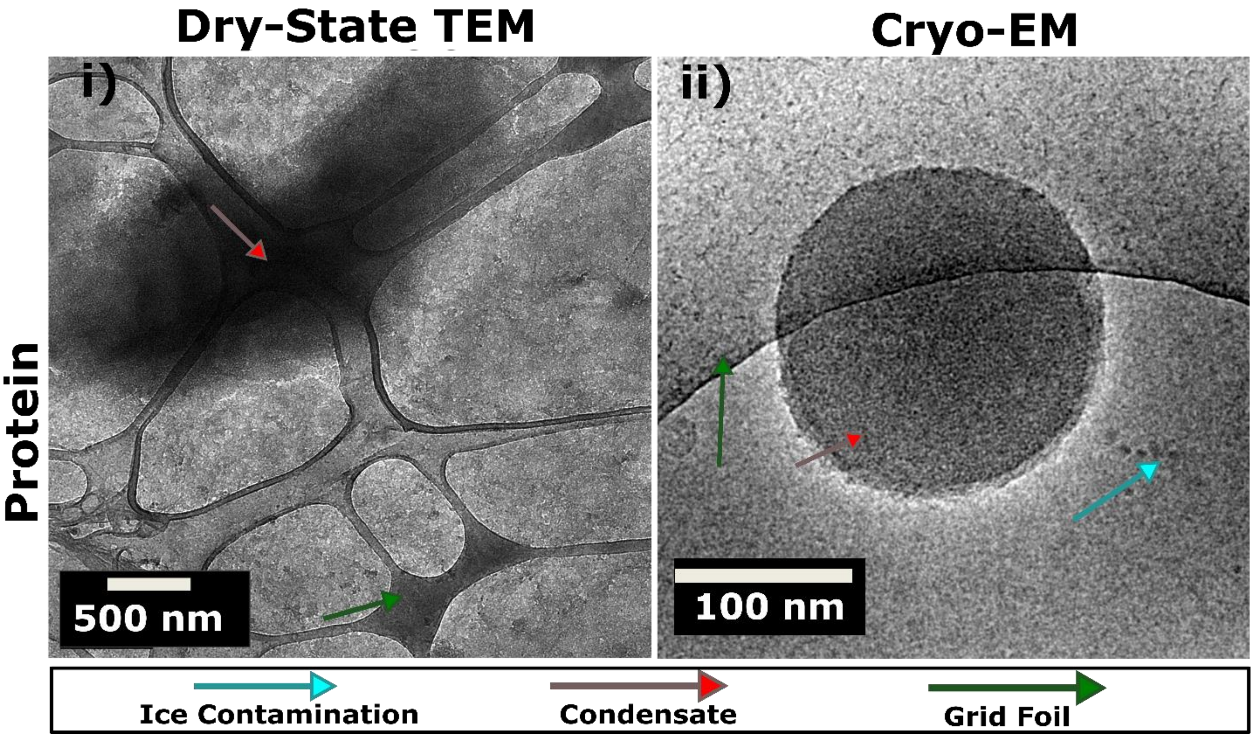


**Supplementary Figure 6:** Cryo-EM vs dry-state comparison of SH condensates i) Dry-state TEM of SH RGG condensate. Dried condensate material can be seen with no preserved structure. ii) Cryo-EM image of SH RGG condensate showing preserved state. Dry-state specimens are prepared on lacey carbon grids which appear as a web-like structure in the background of i.

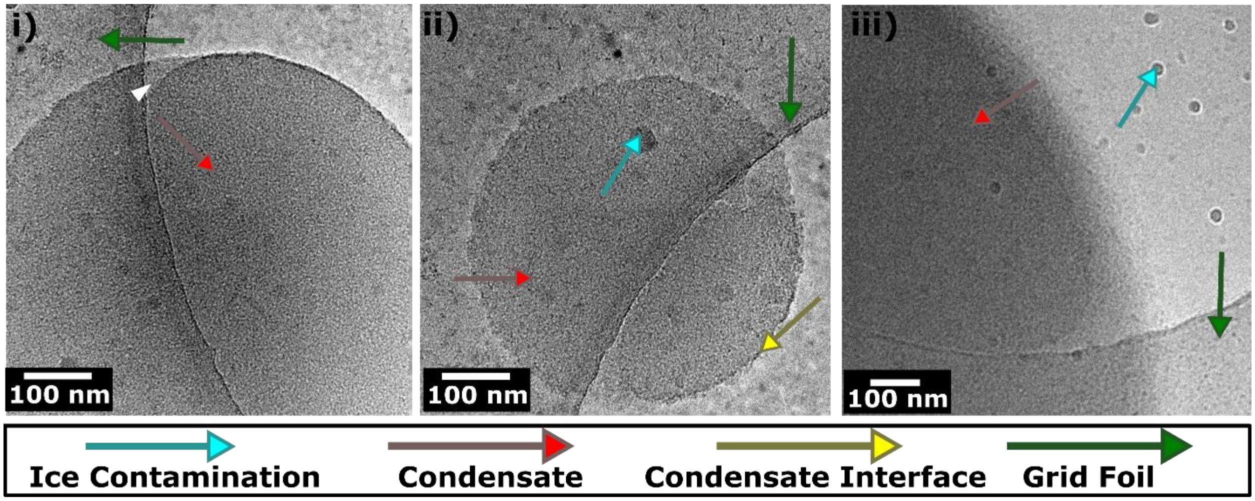


**Supplementary Figure 7:** Irregular interfaces observed during cryoEM imaging. i) ShD condensate with a broken interface. ii) ShD condensate with a patchy interface boundary. iii) ShD WT condensates with a gradient edge and no distinct interface as seen in others.

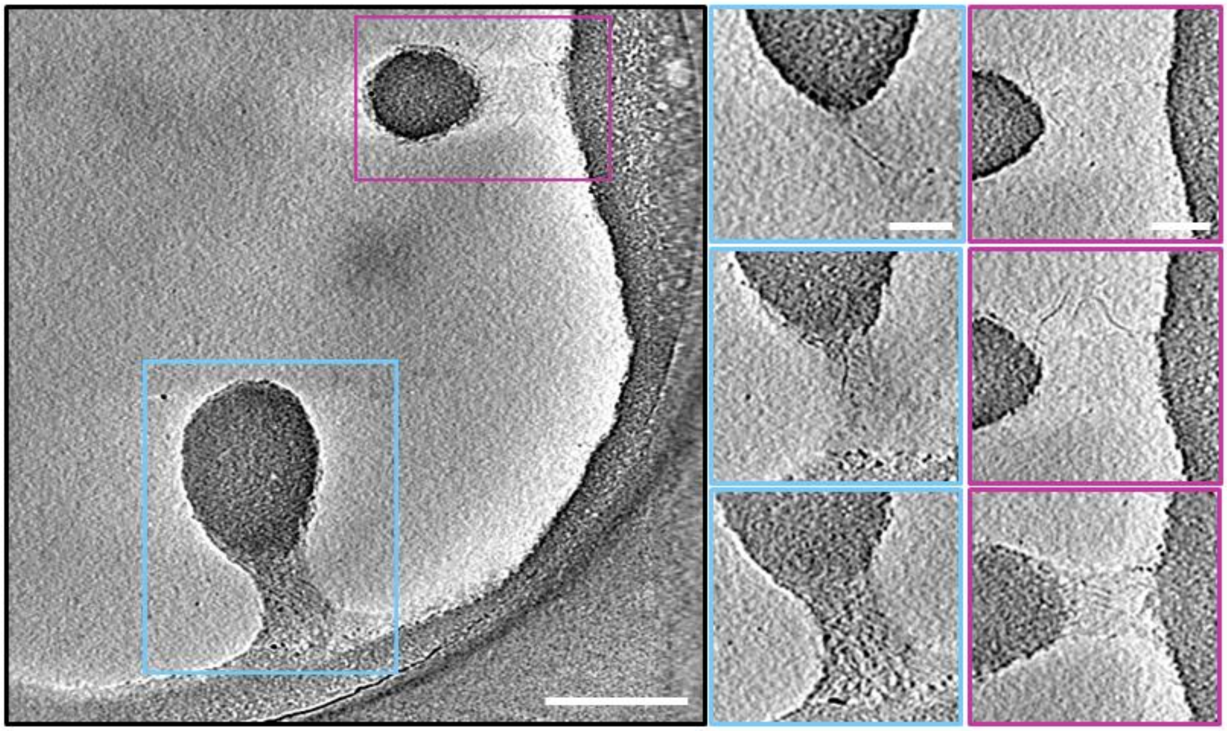

**Supplementary Figure 8:** Examples of RGG WT fiber formation near the liquid-liquid interface. Processed montage of 15 tomographic slices showing two WT condensates pinching off from a larger condensate located on the carbon film. To the right are zoomed in tomographic slices highlighting fibers connecting the smaller condensates to the larger construct. Blue and magenta colors correspond to the condensate boxed in the panel to the left. Fibers were largely observed near smaller droplets that were breaking off from larger condensate material. The breakage process creates a high energy interface and also induces shear forces at the material present at the neck of the droplet breaking off. This supports recent findings showing shear force induced fiber formation in FUS condensates.[7] Furthermore, condensates made from low complexity domains of hnRNPA1 were recently reported to form fibrils at the droplet interfaces.[8] Generally, interfaces are known to promote heterogenous nucleation of fibrils and other aggregates.[9–11] In our work, we only observed such fibers on the nanoscale, suggesting these are metastable morphologies and likely not stable at higher length scales for RGG WT.[12] Scale bar on the large panel is 300 nm and the smaller panel scale bar is 100 nm.

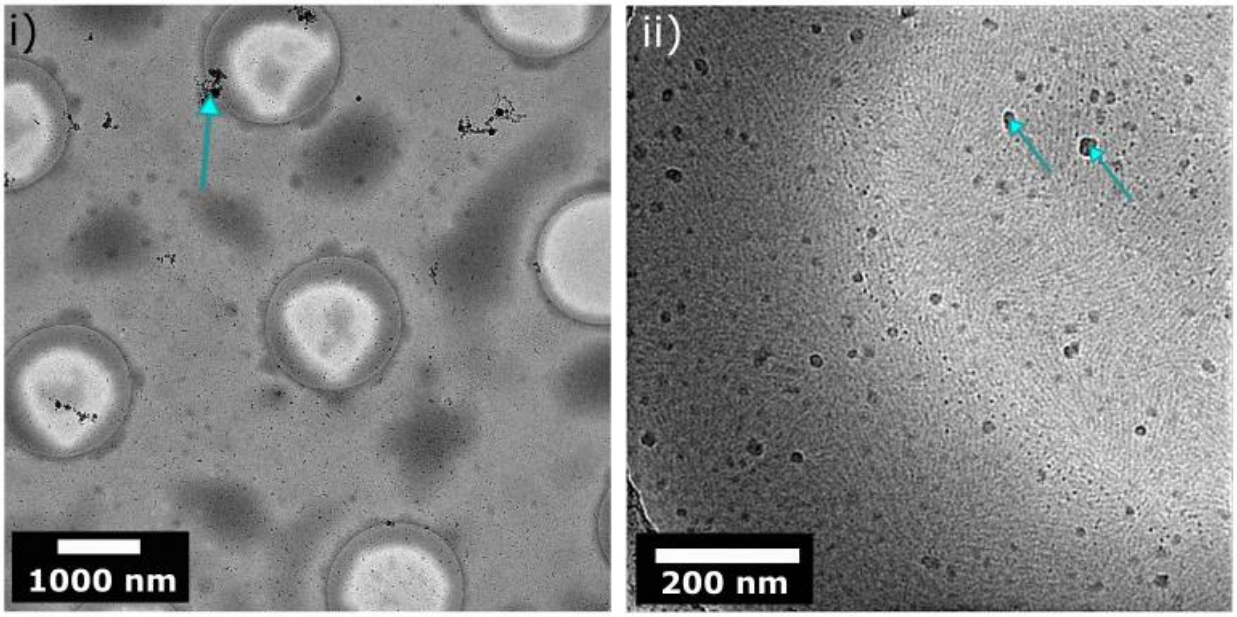


**Supplementary Figure 9:** Shear induced effects on WT RGG. i) Cryo-EM image of WT RGG sample vitrified no post blot wait time. The sample showed no spherical condensates. ii) Closer view of the sample showcasing a shear induced lamellar arrangement of protein. This observation was only made in WT RGG sample. This may be due to the lower viscosity of WT RGG condensates which would mean the condensates can be disturbed more easily as compared to the SH variant.  Blue arrows indicate ice contamination, this sample suffered with a lot of transfer ice.
